# CREBBP lysine acetyltransferase domain mutations create zombie enzymes that alter chromatin loading dynamics and prevent EP300 redundancy

**DOI:** 10.1101/2023.09.05.556242

**Authors:** Haopeng Yang, Wenchao Zhang, Vida Ravanmehr, Ariel Mechaly, Leslie Regad, Ahmed Haouz, Ruidong Chen, Jared M. Henderson, Estela Rojas, Ashley Wilson, Sydney Parsons, Loretta Nastoupil, R. Eric Davis, Qing Deng, Felipe Samaniego, Fernando Rodrigues-Lima, Michael R. Green

**Affiliations:** Department of Lymphoma & Myeloma, University of Texas MD Anderson Cancer Center, Houston, Texas, USA; Université Paris Cité, CNRS, é de Biologie Fonctionnelle et Adaptative Unité de Biologie Fonctionnelle et Adaptative, F-75013 Paris, France; Institut Pasteur, Université Paris Cité, CNRS, Plate-forme de Cristallographie-C2RT, F-75015, Paris, France; Department of Genomic Medicine, University of Texas MD Anderson Cancer Center, Houston, Texas, USA

## Abstract

Epigenetic regulation of gene expression is fundamental to cell state transitions. A prominent example of this is the epigenetically-regulated generation of transitory cells states within the germinal center (GC) reaction that are required for humoral immunity. The deregulation of these processes through somatic mutations can give rise to B-cell lymphoma, thus studying these mutations can provide insight into fundamental mechanisms underlying epigenetic regulation of gene expression as well as GC B-cell and lymphoma biology. Here we show that classes of mutations in the *CREBBP* lysine acetyltransferase gene result in different structure, catalytic activity and function. We discover that CREBBP and its paralog EP300 are dynamically and reciprocally loaded onto chromatin during a signal-responsive cell state transitions within the GC. Mutation of *CREBBP* in GC-derived lymphoma interrupt signal-responsive CREBBP loading onto chromatin, while also inhibiting EP300 redundancy. These observations lead to a model in which CREBBP or EP300 loss of function through acetyltransferase domain mutations, which occur frequently in both hematopoietic and solid tumors, create a zombie enzyme that retards the function of the active paralog by binding limiting concentrations of transcription factor substrate and preventing signal-responsive enhancer activation.

## INTRODUCTION

Cell fate determination requires the activation of signal-responsive epigenetic and transcriptional programs driven by transcription factors and their interacting co-activator or co-inhibitory complexes. This commitment down trajectories of cell fate determination is popularly visualized as the Waddington landscape, in which deterministic junctures constrain subsequent differentiation potential towards defined cell states. In cancer, these processes are perturbed by epigenetic deregulation that biases cell fate determination and/or increases cell plasticity (1,2). Germinal center (GC) B-cell derived malignancies are a prominent example of this due to the complexity and dynamics of the underlying epigenetic programs that regulate normal GC B-cell differentiation, and their deregulation by somatic mutations within genes that encode epigenetic machinery such as histone acetyltransferases and methyltransferases (3).

The germinal center (GC) reaction encompasses two major transitory states in the development and affinity maturation of a B-cell in response to a T-cell dependent antigen: centroblasts and centrocytes. In the dark zone of the GC, centroblasts repress DNA damage sensing, cell cycle checkpoint genes, B-cell receptor (BCR) signaling and antigen presentation through the master transcription factor BCL6 to allow for cells to undergo rapid proliferation and to edit their immunoglobulin genes through somatic hypermutation. Cells then transition into the light zone as centrocytes, where they reinstall programs of BCR signaling and antigen presentation so that they can undergo affinity selection. Centrocytes are multi-potent, able to give rise to centroblasts through dark zone recycling, or either memory B-cells or plasma cells through divergent terminal differentiation programs. These cell fate decisions are made in response to antigen-driven signaling through the BCR and through signaling from other cells within the light zone of the GC, including CD40 ligand and cytokines from T follicular helper (T_FH_) cells. Interaction with T_FH_ cells is competitive, and the strength of CD40 signaling received directly impacts the cell fate decisions towards recycling or terminal differentiation. These signaling-responsive cell state transitions are manifested through the activation of transcription factors and subsequent chromatin remodeling that alters the enhancer activity and expression of thousands of genes (4). In follicular lymphoma (FL) and the GC B-cell subtype of diffuse large B-cell lymphoma (GCB-DLBCL), the dynamics of epigenetic programs required for GC B-cell state transitions are perturbed via chromatin modifying gene (CMG) mutations (5). The *CREBBP* (a.k.a. Cbp) gene is the second most frequently mutated CMG in FL and GCB-DLBCL and, together with its paralog *EP300* (a.k.a. p300), encode a lysine acetyltransferase that is recruited to chromatin by transcription factors to act as a transcriptional co-activator via acetylation of histone 3 lysine 27 (H3K27) and other residues. Mutations of *CREBBP* most commonly occur as missense mutations within the catalytic lysine acetyltransferase (KAT) domain and have been described as being loss of function (6,7). CREBBP and EP300 are paralogous enzymes that are thought to be largely redundant, to the extent that they are often collectively described as Cbp/p300. However, studies using compound conditional knock-out of *Crebbp* and *Ep300* within murine GC B-cells have suggested that they may have some non-redundant functions (8). In addition, the most frequent mutation of *CREBBP* (R1446C) has a more pronounced phenotype than *CREBBP* knock-out (KO) (9), suggesting that there may be effects that extend beyond simple loss-of-function. However, the mechanism for these effects, whether it is true for other mutations, and the precise roles for CREBBP and EP300 in normal GC B-cells and their malignant counterparts remain to be defined.

Here we show using crystal structure and biochemical analysis that common hotspot mutations within the *CREBBP* KAT domain can have divergent effects on the acetyl-CoA binding pocket, resulting in classes of mutations with different catalytic activity. Using CRISPR editing of lymphoma cell lines and integrative epigenetic profiling, we show that these mutation classes have respective differences in the expression of cytokine-and NFκB-responsive gene sets, but maintain a core program of epigenetically-repressed genes that are characteristic of centrocyte gene expression programs. *CREBBP* KAT domain missense mutations were universally able to inhibit the redundant activity of EP300 despite ineffective chromatin loading, suggesting the interruption of relative dynamics. Through analysis of human GC B-cells, we discover that CREBBP and EP300 are dynamically and reciprocally loaded or unloaded from chromatin during the centroblast to centrocyte transition, respectively. Using functional analysis we show that CREBBP is dynamically loaded onto chromatin in response to CD40 signaling, which is impaired by the loss of catalytic activity. Furthermore, we show that mutant CREBBP prevents the redundant activity of its paralog EP300 by binding limiting quantities of transcription factor, akin to zombie enzymes that are encoded by catalytically inactive pseudogenes that alter the function of their active paralogs (9). This therefore identifies a new mechanism of oncogenesis via perturbed signal-responsive cell state transitions resulting from mutationally-encoded zombie enzymes, which likely applies to both hematopoietic and solid tumors that acquire *CREBBP* or *EP300* KAT domain mutations.

## RESULTS

### *CREBBP* mutational distribution differs significantly between diffuse large B-cell lymphoma (DLBCL) and follicular lymphoma (FL)

Mutations of *CREBBP* and *EP300* are highly recurrent in both solid tumors and hematological cancers (**Figure S1A-G**) (10,11). However, hematological cancers have a strong bias towards mutating *CREBBP* compared to *EP300*, particularly within non-Hodgkin lymphomas, suggesting a unique function for *CREBBP* in these primarily GC B-cell-derived malignancies. We therefore curated mutation data from 1,028 FL, 188 transformed FL (tFL) and 3,104 DLBCL tumors to characterize the spectrum of *CREBBP* mutations, which were identified in 60.4% of FL (621/1028), 48.4% of tFL (91/188) and 13.8% of DLBCL (427/3104) tumors (Table S1-2) (12–22). The majority of mutations were missense changes within the catalytic KAT domain (782/1380; **Figures 1A** and S1C-D), with the most recurrent mutations being clustered around the acetyl-CoA binding pocket (**Figure 1B**), including hotspots at R1446 (187/1380), Y1503 (94/1380), Y1482 (43/1380) and W1502 (41/1380). An additional hotspot consisting of a single amino acid deletion was observed outside the catalytic pocket at S1680 (69/1380). Nonsense/frameshift mutations were observed in 301 tumors and most often occurred upstream to, or within, the KAT domain (280/301). We observed a significant difference in the relative frequency of mutation types between FL and DLBCL (**Figure 1C**), with FL tumors significantly more frequently harboring R1446 mutations (18.9% of all FL *CREBBP* mutations, found in 13.8% of all FL tumors; Chi-square P-value < 0.001) and DLBCL significantly more frequently harboring nonsense/frameshift mutations (34.6% of all *CREBBP* mutations in DLBCL, found in 5.3% of DLBCL tumors). The higher frequency of *CREBBP* mutations compared to *EP300*, and differential *CREBBP* mutational spectrum between FL and DLBCL therefore suggest a potentially unique role for CREBBP and hint at important functional differences between classes (KAT missense vs nonsense/frameshift) and sites (eg. R1446 vs Y1482/Y1503) of mutation.

**Figure 1:**
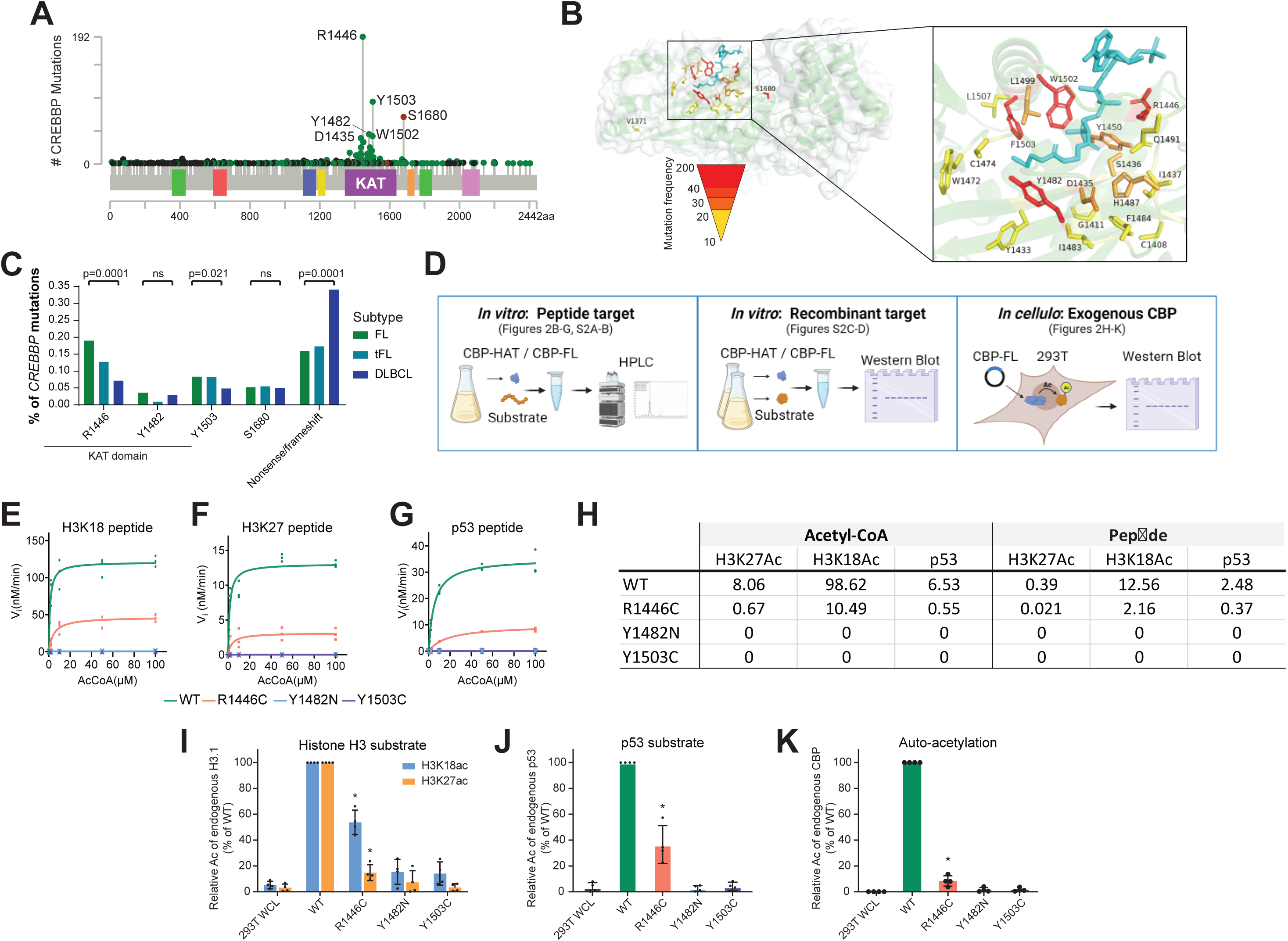
Frequency and biochemical characteristics of recurrent *CREBBP* lysine acetyltransferase (KAT) domain mutations. **A)** A lollipop plot of 1,139 CREBBP mutations curated from 4,320 lymphomas showing hotspots of missense mutations within the KAT domain at R1446, Y1503 and Y1482. **B)** Rendering of recurrent (observed ≥10 times) KAT domain missense mutations over the CREBBP structure, colored by frequency, showing their relationship to the acetyl-CoA binding pocket. **C)** The frequency of CREBBP hotspot mutations and nonsense/frameshift (truncating) mutations in FL, tFL and DLBCL. **D)** A schematic of the orthogonal approaches taken to evaluate the activity of *CREBBP* KAT domain mutants. **E-G)** Vi determined by RP-UFLC for (E) H3K18, (F) H3K27 and (G) p53 peptides across varying concentrations of Acetyl-CoA. **H)** Vmax/Km values for WT and KAT point mutant CREBBP determined by RP-UFLC with varying concentrations of Acetyl-CoA or target peptide. **I-K)** *In cellulo* assessment of the activity of full-length CREBBP by transfection into 293T cells and western blot assessment of endogenous (I) histone H3, (J) p53 or (K) CREBBP.

### KAT domain hotspot mutations of *CREBBP* have distinct biochemical consequences

We hypothesized that mutations with differential representation across lymphomas may be selected for or against due to unique functional consequences on CREBBP catalytic activity. We explored this using acetyltransferase assays with WT, R1446C, Y1482N and Y1503C recombinant human full-KAT (CREBBP-KAT: catalytic core that includes Bromo, Ring, PHD and KAT domain) or transfected full-length human protein (CREBBP-FL) with multiple orthogonal approaches (**Figure 1D**). We measured *in vitro* acetyltransferase activity by reverse-phase ultra-fast liquid chromatography (RP-UFLC) (23) using (i) recombinant CREBBP-KAT with a fixed concentration of 30µM peptide substrate (H3K18, H3K27, p53) and increasing concentration of acetyl-CoA (**Figures 1E-G**); (ii) CREBBP-KAT with a fixed concentration of 100µM acetyl-CoA and increasing concentrations of peptide substrate (**Figure 1H-J**); or (iii) end-point assays with CREBBP-KAT or immunoprecipitated CREBBP-FL and fixed concentrations of 100µM acetyl-CoA and 30µM substrate (Figures S1E-F). All assays showed a highly reproducible loss of activity for R1446C compared to WT, and no detectable activity above baseline for Y1482N or Y1503C mutants. Specifically, the catalytic efficiency of R1446C (Vmax/Km) was 5.4-17.2% lower than the WT protein, depending on the assay (**Figure 1E-H**). These trends of reduced activity for R1446C and undetectable/background activity of Y1482N and Y1503C were also observed using *in vitro* acetyltransferase assays with standardized concentrations of acetyl-CoA and recombinant protein substrates (histone H3.1/2/3 or p53) with either CREBBP-KAT (Figure S1H-I) or CREBBP-FL (**Figure 1I-J**). However, these assays additionally revealed substrate-specific differences in loss of catalytic activity for the R1446C mutant. Specifically, the activity of R1446C is less impaired when employing H3K18 as a substrate compared to when using H3K27 as a substrate, in line with a preference of CREBBP for H3K18 as a substrate (24). CREBBP is also able to catalyze autoacetylation, including on lysines within its autoinhibitory loop (25). We assessed this in 293T cells transfected with CREBBP-FL by immunoprecipitation and western blot for acetyl-lysine (AcK). The R1446C mutant retained low but detectable autoacetylation activity, but no detectable autoacetylation above the background of the assay was observed for the Y1482N and Y1503C mutants (**Figure 1K**). Lower but detectable levels of acetylation were also observed with an AIL-deleted mutant of full-length CREBBP (Figure S1K) indicating that autoacetylation can occur outside of these residues. These biochemical assays therefore revealed differences in catalytic activity between highly-recurrent *CREBBP* mutations, with the most frequent mutation (R1446C) partially impairing its catalytic activity in a substrate-specific manner, and other less frequent but highly recurrent mutations (Y1482N and Y1503C) resulting in loss of any detectable catalytic activity.

### CREBBP KAT domain hotspot mutations differentially affect acetyl-CoA interaction

Having observed that the most highly recurrent mutations of *CREBBP* cluster around the acetyl-CoA binding pocket, and have differential activity in biochemical assays, we hypothesized that these mutations may have differential effects on the structural properties of CREBBP and substrate interactions. The KAT domain of CREBBP is highly conserved between mouse and human, including the R1446, Y1482 and Y1503 residues (**Figure 2A**). Crystallization requires complete removal of acetylation and highly flexible regions, and cannot be achieved for a fully WT protein. Prior structural studies of CREBBP and EP300 have addressed this by introduction of an inactivating Y1503F or Y1467F mutations, respectively, into the murine full-KAT (amino acids 1079-1751, incorporating the bromodomain, ring domain, PHD finger, KAT domain and ZZ domain), in addition to removal of the flexible AIL from the protein (26–29). We therefore followed this same approach using a murine full-KAT of Crebbp. The Y1503F mutation is itself a frequent mutation observed in lymphoma (Table S3) and maintains the aromatic ring but removes the hydroxyl group. We additionally generated a Y1503C variant, which we found to be catalytically inactive and which leads to loss of the aromatic ring at this position. The Y1482N mutation was also found to be catalytically inactive and therefore did not have to be combined with the Y1503F mutation, allowing assessment of the WT configuration of Y1503 within the Y1482N structure. However, because the R1446C mutation maintained detectable activity in biochemical assays, it was combined with the Y1503F mutation to eliminate its catalytic activity and make it amenable to crystalization. We therefore generated a suite of four variant full-KATs (Y1503F, Y1503C, Y1482N, and R1446C[+Y1503F]) for structural analysis of recurrent KAT domain mutations within *CREBBP* (**Figure 2A**), with Y1482N serving as the control for Y1503F/Y1503C, and Y1503F serving as the control for Y1482N and R1446C.

**Figure 2:**
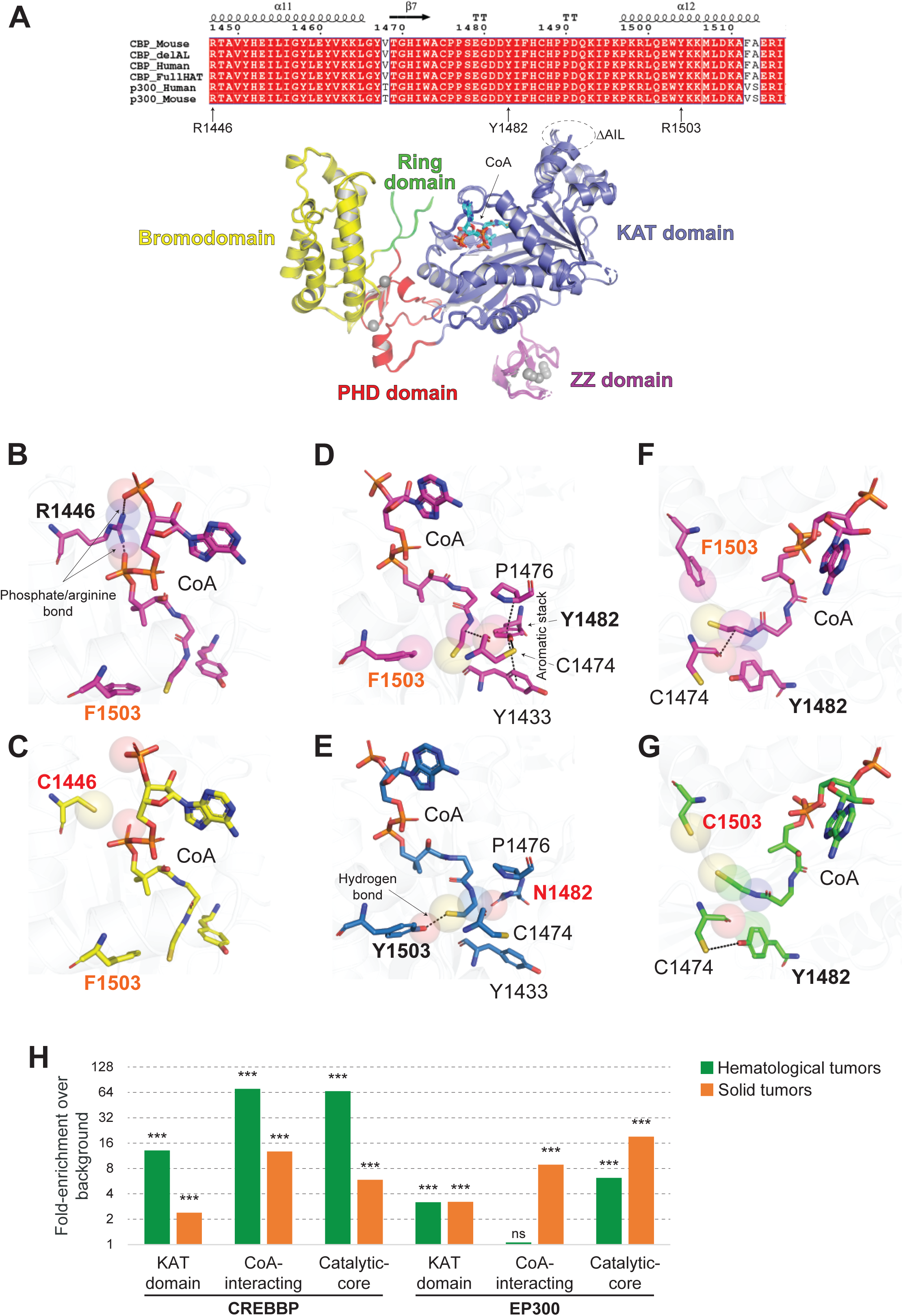
Crystal structure analysis of CREBBP KAT domain mutants. **A)** Alignment of the human and mouse KAT domains over the area including the mutations, and an overview of the crystal structure. **B-C)** Focused evaluation of R1446 (WT) and C1446 (mutant) in the F1503 mutant background showing a loss of interaction with CoA. **D-E)** Evaluation of Y1482 (WT) in the F1503 mutant background, and N1482 with Y1503 WT background. The loss of an aromatic stack, positioning of C1474 and enlargement of the catalytic core can be observed with N1482 mutation. The Y1503 WT allele can be seen to form a hydrogen bond with the β-mercapto-ethylamine moiety of CoA. **F-G)** Evaluation of F1503 and C1503C mutants showing a loss of interaction with the β-mercapto-ethylamine moiety of CoA and enlargement of the catalytic core. H) Analysis of COSMIC mutation data for hematological (green) and solid (orange) tumors showing enrichment of missense mutations within the KAT domain, and targeting residues that are predicted from crystal structure analysis to interact directly with the distal portion of CoA or to shape the catalytic core. ***Fisher exact test P-value < 0.0001.

High purity, deacetylated protein was produced, co-crystalized with coenzyme-A (CoA), and analyzed by x-ray crystallography to produce structures with 2.466lJ (Y1503F), 2.016lJ (Y1503C), 2.450lJ (Y1482N) and 2.287lJ (R1446C+Y1503F) resolution (Table S4). The overlay of the structures presented in Figure 2A showed good agreement with the previously published structure of CREBBP in apo form (PDB ID: 5U7G; Figure S2A) (28).

The only observable differences between structures were found in the interaction between CREBBP and CoA, thus these interactions will be the focus. Evaluation of the R1446 residue within the Y1503F structure highlighted the special role of this highly-mutated amino acid in forming a strong phosphate/arginine salt bridge with the pyrophosphate and adenosine phosphate moieties of CoA (**Figure 2B**). The R1446C mutation eliminates this interaction and enlarges the cofactor binding pocket resulting in slight movement of the pantothenic acid and adenosine diphosphate moieties, but maintains the position of the β-mercapto-ethylamine moiety within the catalytic core of the enzyme (**Figure 2C** and Figure S2B). By evaluating the Y1482 position within the Y1503F structure, we observed that Y1482 forms an aromatic stack with Y1433 and P1476 to create a hydrophobic substrate binding tunnel in which C1474 is positioned to form a hydrogen bond with the β-mercapto-ethylamine group of CoA (**Figure 2D**). The Y1482N mutation disrupts the aromatic stack, slightly enlarging the substrate binding tunnel, and alters its hydrophobicity (Figure S2C), leading to a change in positioning of the β-mercapto-ethylamine group in the catalytic core of the enzyme (**Figure 2E**). Further biochemical analysis of amino acid variants at this position revealed that Y1482F (hydrophobic, aromatic ring) but not Y1482H (hydrophilic, aromatic ring) or Y1482L (hydrophobic, no aromatic ring) maintained acetyltransferase activity *in vitro* (Figure S2D), indicating that both hydrophobicity and maintenance of the aromatic stack is required for catalytic activity. Close inspection of Y1503 within the Y1482N structure revealed a hydrogen bond between the hydroxyl group of Y1503 and the terminal sulfur atom of the β-mercapto-ethylamine moiety of CoA from which the acetyl group is donated (**Figure 2E**). This H-bond has been suggested to be important for catalysis for p300 (26) and was eliminated in the Y1503F (loss of the hydroxyl group of the side chain) (**Figure 2F**) and Y1503C (loss of the hydroxyl group and less bulky side chain) (**Figure 2G**) mutants contributing to an enlargement in the catalytic core of the enzyme, and a change in the positioning of the β-mercapto-ethylamine moiety within the catalytic core was observed for Y1503C. Together our biochemical and structural data indicate that mutations that alter the positioning of the β-mercapto-ethylamine moiety, and therefore likely the acetyl group, within the catalytic core of CREBBP (Y1503F/C, Y1482N) lead to a loss of catalytic activity. Other mutations such as R1446C that alter interactions between CREBBP and other moieties of CoA without affecting positioning of the β-mercapto-ethylamine moiety in the catalytic core lead to a reduction in catalytic activity. Using this information, we created a map of *CREBBP* and *EP300* mutations across hematopoietic and solid tumors from COSMIC data (10) that classed variants into (i) truncating mutations within or upstream from the KAT domain, (ii) other truncating mutations, (iii) KAT domain CoA-interacting missense mutations that are likely to reduce activity, (iv) KAT domain catalytic core missense mutations that are likely to be catalytically dead, (v) missense mutations of indeterminate significance inside the KAT domain, and (vi) missense mutations of indeterminate significance outside the KAT domain (Figure S1A-B). Notably, for both genes (*CREBBP* and *EP300*) and in both of the broad groups of malignancies (hematological and solid tumors), missense mutations were 2.6-to 3.7-fold more frequent than truncating mutations. Furthermore, missense mutations were significantly enriched within the KAT domain and at amino acids predicted to interact with CoA or reside within the catalytic core compared to that expected by chance, with the sole exception of EP300 mutations for CoA-interacting residues in hematological tumors (**Figure 2H**). Therefore, although the greatest enrichment was seen for CoA-interacting amino acids (70.6-fold higher than that expected by chance) and amino acids within the catalytic core (66.2-fold higher than that expected by chance) in hematological malignancies, the structural and biochemical principles that we describe here for *CREBBP* mutations in lymphoma can likely be also extrapolated to solid tumors, and in addition likely also apply to the *EP300* gene.

### Catalytically divergent mutations have common and unique epigenetic consequences

The divergent structural and biochemical properties of different *CREBBP* hotspot mutations led us to hypothesize that they may result in different epigenetic phenotypes and transcriptional activity. We therefore extended upon our previous set of CRISPR-engineered lymphoma cell lines (9) to produce isogenic WT, homozygous *CREBBP* knock-out (KO), R1446C, Y1482N and Y1503C mutants in two B-cell lymphoma cell lines (RL and HT) and in 293T cells by editing the endogenous coding sequences (Figure S3A). The RL and HT cell lines have monoallelic frameshift mutations within the *EP300* gene, allowing for less confounded assessment of the effects of changes in CREBBP catalytic activity, and 293T cells are WT for EP300. All selected clones with point mutations introduced into *CREBBP* were homozygous, which is observed for ∼40% of CREBBP mutations in primary FL tumors (12), and mutations did not change the expression of CREBBP or EP300 at the protein level (Figure S3B). We molecularly profiled these isogenic lines by CUT&RUN (29) for H3K27Ac, H3K4me1, H3K4me3, H3K27me3, CREBBP, EP300 and BRD4; assay for transposase-accessible chromatin (ATAC)-sequencing; and RNA-sequencing. All consensus H3K27Ac peaks between RL and HT WT cells were considered, and classified as super-enhancers by ROSE (30) (Figure S3C), enhancers by an overlapping H3K4me1 peak and no overlapping H3K4me3 peak, promoters by an overlapping H3K4me3 peak, and weak enhancers with low H3K27Ac and no overlapping H3K4me1 or H3K4me3 peak (Figure S3D). CREBBP CUT&RUN was validated against previously published ChIP-seq and showed good concordance (Figure S4), but a minor level of background signal was observed in CREBBP KO cells likely as a consequence of some recognition of EP300 by the polyclonal antibody. Technical duplicates were performed for each assay, then the data merged for each clone. For this analysis, we utilized an H3K27Ac consensus peak set for RL and HT lymphoma cell lines, agnostic to CREBBP or EP300 loading, and analyzed 293T cells separately (Figure S5). Unsupervised principal component analysis (PCA) of H3K27Ac and RNA-seq revealed that KO clones were more similar to WT than any of the point mutant clones, and R1446C mutants (partial loss of function) were reproducibly different to Y1482N and Y1503C clones at the epigenetic and transcriptional level (**Figure 3A-B**, Figure S5). As biochemical analysis revealed that these mutations resulted in a loss of catalytic activity, we focused on H3K27Ac loss, and catalytically dead mutants (Y1482N and Y1503C) were considered collectively due to their biochemical and molecular similarities. Differential analysis of H3K27Ac between R1446C or Y1482N/Y1503C mutants and WT cells identified regions unique to, or shared by, each class of KAT mutation (**Figure 3C-D**; Table S6). This pattern was reproducible within the 293T cell line (Figure S5). The most notable difference between mutational classes were the distribution of the R1446C-or Y1482N/Y1503C-specific H3K27Ac loss regions according to their genomic annotation (**Figure 3E**). H3K27Ac loss most frequently affected enhancers, but Y1482N/Y1503C-specific regions significantly more frequently affected super-enhancers (significant H3K27Ac loss in 70% (802/1138) of super-enhancers; Chi-square P-value = 0.0001) and weak enhancers (significant H3K27Ac loss in 12% (778/6621) of weak enhancers; Chi-square P-value = 0.0001) (**Figure 3E**). Thus, the low level of catalytic activity in R1446C mutants may be sufficient to maintain weak enhancers and super-enhancers resulting in 79% (1145/1453) of R1446C-specific H3K27Ac loss regions affecting generic enhancers. In contrast, although a higher number of enhancers showed loss of H3K27Ac in Y1482N/Y1503C-specific regions (n=1498), the lack of catalytic activity may have a greater effect on weak enhancers and particularly on super-enhancers, with a loss of H3K27Ac being observed in 83% of super-enhancers (946/1138) when considering both Y1482N/Y1503C-specific and shared regions. Gene annotation of these regions (closest gene) and hypergeometric gene set enrichment analysis (hGSEA) of B-cell and lymphoma-related gene sets showed a modest enrichment of pathways associated with cytokine response in R1446C-specific loss regions and NFκB target gene sets in Y1482N/Y1503C-specific loss regions (**Figures 3F** and Table S7). The most significant enrichment was noted among regions found to have H3K27Ac loss in both the R1446C-vs-WT and Y1482N/Y1503C-vs-WT comparisons, highlighting a significant enrichment of major histocompatibility complex (MHC) genes, centrocyte (CC) signatures, CD40-responsive gene sets, and gene repressed by BCL6. Biochemical differences between classes of *CREBBP* KAT domain mutations may therefore have relatively different impact upon cytokine response or NFκB activity, but the core set of commonly deregulated regions are more strongly enriched for pathways associated with lymphoma pathobiology including the cell state transition between centroblasts (CB) and CC, and associated immune synapse formation and signaling response.

**Figure 3:**
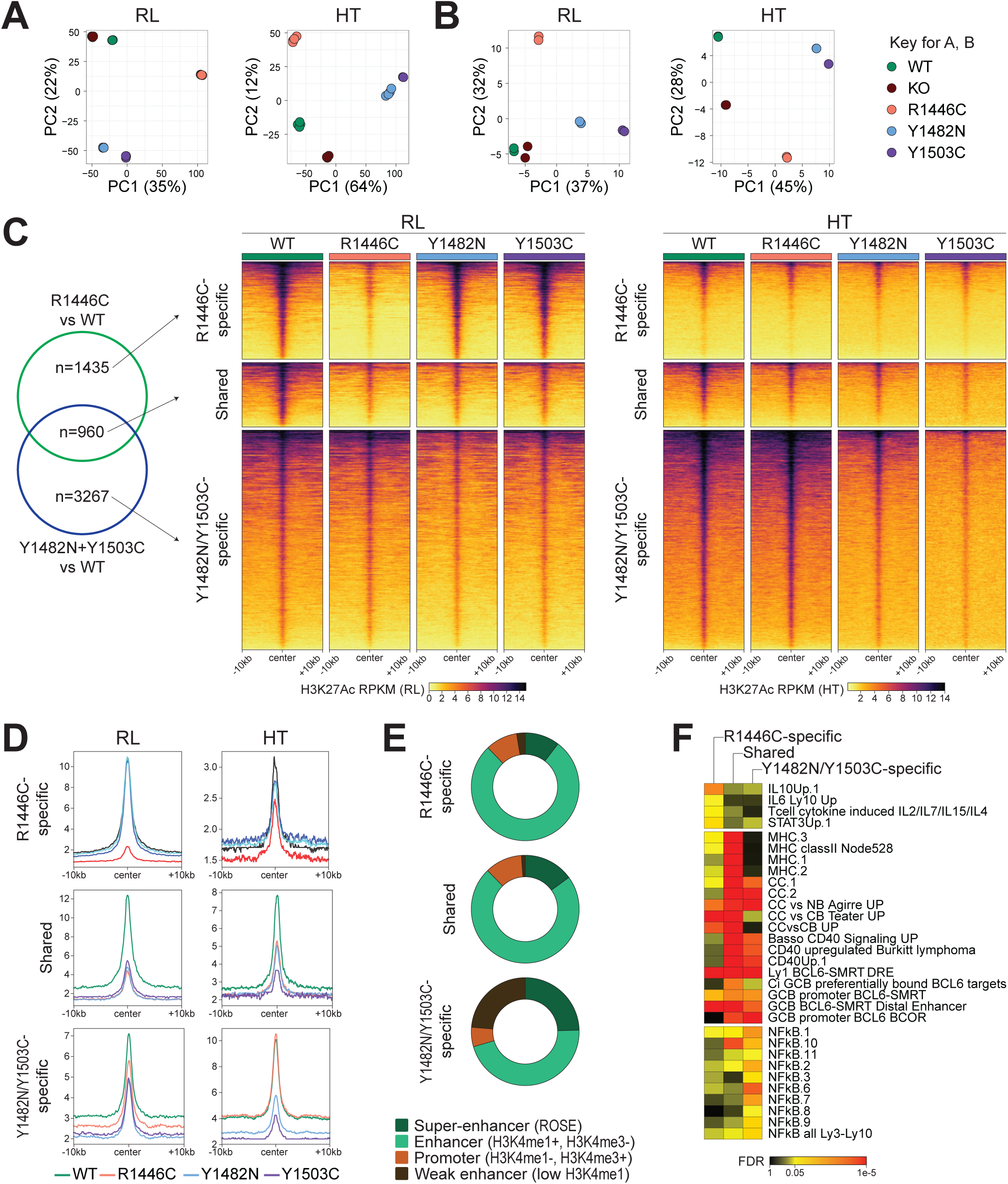
Molecular differences between R1446C and Y1482N/Y1503C mutants. **A-B)** Unsupervised principal component analysis (PCA) plots for (A) H3K27Ac CUT&RUN and (B) RNA-sequencing showing divergence of KAT mutants from WT and KO cells, and differences between catalytically dead mutants (Y1482N and Y1503C) and R1446C. **C-D)** Supervised analysis of H3K27Ac CUT&RUN comparing R1446C or Y1482N+Y1503C to WT cells identifies R1446C-specific, shared and Y1482N/Y1503C-specific regions of H3K27Ac loss. **E)** Distribution of H3K27Ac loss regions within each category from (C) according to genomic annotation showing an over-representation of super-enhancers and weak enhancers within Y1482N/Y1503C-specific regions. **F)** Hypergeometric gene set enrichment analysis of genes associated with R1446C-specific, shared and Y1482N/Y1503C-specific regions.

### KAT domain point mutations (KAT-PM) reduce CREBBP chromatin loading and prevent the redundant activity of EP300

Unsupervised analysis of H3K27Ac and RNA-seq showed that biallelic *CREBBP* KO maintained a phenotype more similar to WT cells than either class of KAT domain point mutants (KAT-PM), suggesting a potential compensatory role for EP300 in the setting of *CREBBP* KO that is absent in the setting of KAT-PM. Given the shared enrichment of important GC B-cell programs between different KAT-PMs, we investigated this by defining the consensus regions of H3K27Ac loss between WT and KAT-PM (R1446C/Y1482N/Y1503C) across both lymphoma cell lines (RL and HT). In order to enrich the likelihood of observing CREBBP/EP300 redundancy, we focused on H3K27Ac peaks with overlapping CREBBP and EP300 peaks shared between both RL and HT WT cells (n=26,164). Differential analysis between WT and KAT-PM cells identified 2,302 regions of significant H3K27Ac loss (FDR<0.05, fold-change>1.5; **Figure 4A**; Table S8) corresponding to super-enhancers (n=504 peaks within 319), enhancers (n=1462) and promoters (n=521). Loss of H3K27Ac in KAT-PM cells was associated with loss of H3K4me1 in super-enhancers and enhancers, loss of H3K4me3 in promoters, loss of BRD4 loading across all regions (**Figure 4B**). We also observed an increase in H3K27me3 at regions of H3K27Ac loss (**Figure 4C**), suggestive of enhancer/promoter decommissioning. However, we observed a minimal loss of chromatin accessibility (**Figure 4B**). In *CREBBP* KO cells we noted similar levels as WT cells or a markedly-lower magnitude of reduction of activating marks (**Figure 4A-B**) and less gain in H3K27me3 (**Figure 4C**) relative to that observed in KAT-PM cells. This provides further support for a potential compensatory role of EP300 in *CREBBP* KO cells that is absent *CREBBP* KAT-PM cells. We confirmed patterns of EP300 compensation by evaluating its loading by CUT&RUN, which showed increased loading in *CREBBP* KO cells but not in *CREBBP* KAT-PM cells (**Figure 4D-E**). We hypothesized that this may be due to the presence of a catalytically-inactive CREBBP protein at these sites in KAT-PM cells, however analysis of CREBBP CUT&RUN showed a significant reduction in CREBBP loading in KAT-PM cells over super-enhancers, enhancers and promoters with loss of H3K27Ac (**Figure 4F-G**). Therefore, loss of catalytic activity in *CREBBP* KAT-PMs results in reduced chromatin loading together with concomitant failure of EP300 to compensate at these loci, resulting in enhancer and promoter decommissioning.

**Figure 4:**
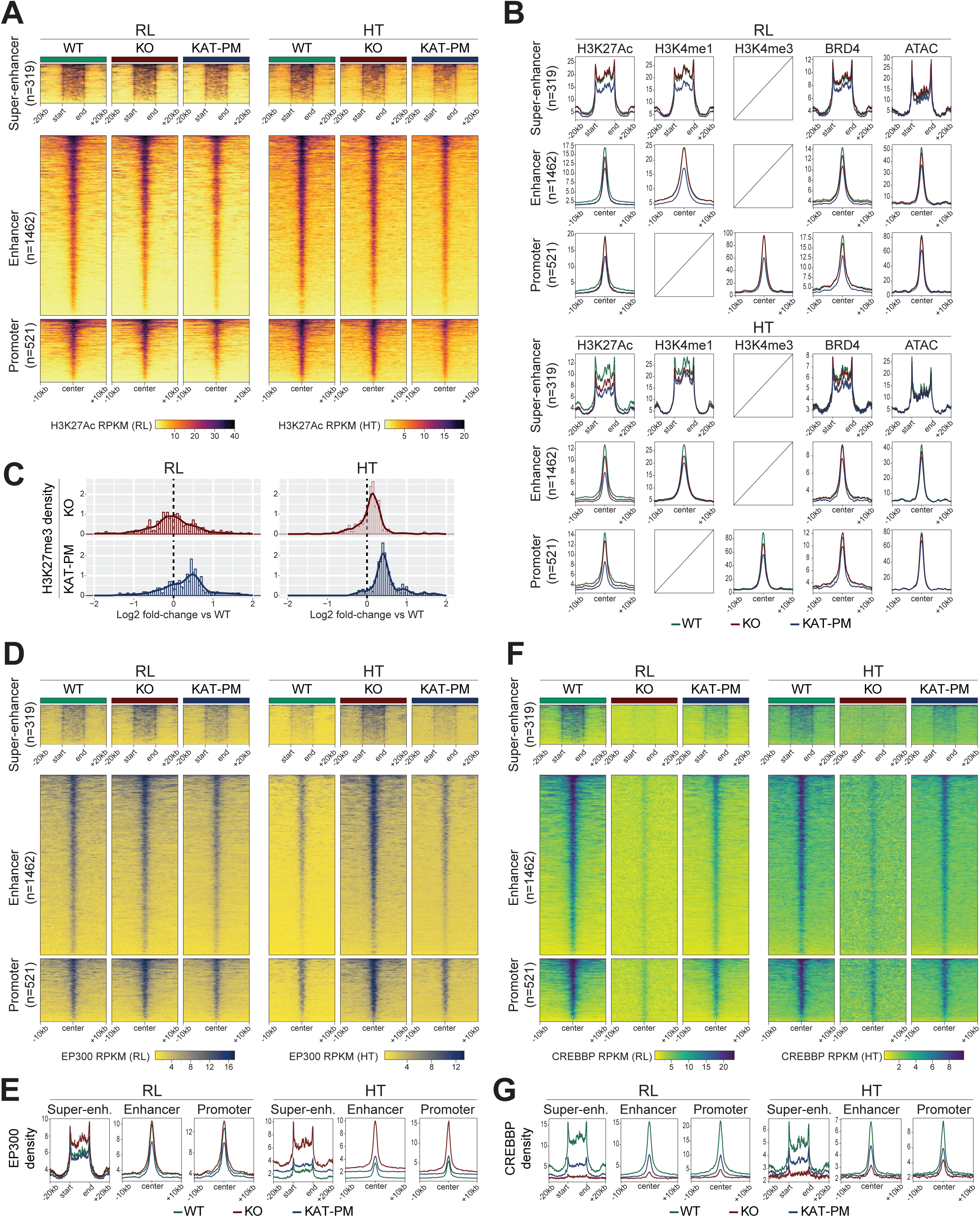
Analysis of KAT domain point mutants (KAT-PM) compared to biallelic knock-out (KO). **A)** Heatmaps show super-enhancers, enhancers and promoters with significant (FDR≤0.05, fold-change≥1.5) loss of H3K27Ac in KAT-PM compared to WT, with KO shown for reference. **B)** Density plots of H3K27Ac, H3K4me1 (active enhancer mark), H3K4me3, BRD4 and ATAC (chromatin accessibility) across regions shown in (A) for the RL (above) and HT (below) isogenic cell lines. **C)** Log2 fold-change of H3K27me3 reads within peaks shown in (A) comparing KO (above) or KAT-PM (below) to isogenic WT cells for RL (left) or HT (right). **D-E)** Heatmaps (D) and density plots (E) for EP300 loading over regions with H3K27Ac loss in KAT-PM cells compared to isogenic WT cells, as shown in (A). **F-G)** Heatmaps (D) and density plots (E) for CREBBP loading over regions with H3K27Ac loss in KAT-PM cells compared to isogenic WT cells, as shown in (A).

### *CREBBP* mutations alter normal dynamics of CREBBP loading in the germinal center reaction

Having observed that *CREBBP* KAT-PMs result in a loss of promoter/enhancer activity and CREBBP loading at gene set involved in the CB to CC cell state transition, we reasoned that *CREBBP* mutations may deregulate a normal dynamic processes of CREBBP loading that occurs during this process. We first investigated this by sorting highly-purified CBs and CCs from 3 human tonsils (**Figure 5A**) and evaluating patterns of CREBBP and EP300 loading by CUT&RUN. Differential analysis identified thousands of loci with significant (FDR<0.05, fold-change>1.5) changes in CREBBP and EP300 loading between CBs and CCs (**Figure 5B-C**). This included a predominant pattern of CREBBP loading (CB < CC; 2589/3858 significant regions) and EP300 unloading (CB > CC; 2530/2772 significant regions) in CCs compared to CBs (**Figure 5B-C**). These dynamically loaded regions were largely non-overlapping (**Figure** 5D), consistent with non-redundant roles for EP300 and CREBBP in CBs and CCs, respectively (7). Notably, when comparing WT and KAT-PM lymphoma cells we observed a significant reduction of CREBBP loading over the same regions that are dynamically loaded with CREBBP in the non-malignant CB to CC transition (**Figure 5E-F**). This is consistent with a failure of CREBBP KAT-PM to be loaded onto chromatin at these developmentally-regulated loci. hGSEA revealed further similarities between regions that are dynamically loaded with CREBBP in the CB to CC transition and regions with a loss of CREBBP loading in KAT-PM (**Figure 5G**; Table S9). Specifically, these gene sets were significantly enriched for genes with reduced expression in primary follicular lymphomas with CREBBP mutations (12), MHC gene sets, BCL6-repressed genes, CC signature genes, and genes that are responsive to CD40 signaling including IRF4 and NFκB target genes (**Figure 5G**). We further explored this via motif enrichment analysis, which showed selective enrichment of motifs for CD40-responsive AP-1, NFκB and IRF transcription factor families within regions loaded with CREBBP in the CB to CC transition and those with CREBBP unloading in KAT-PM cells (**Figure 5H**; Table S10). These regions included genes involved in antigen presentation (e.g. *HLA-DRA* and *CD74*) and memory B-cell differentiation (e.g. *BACH2*) (**Figure 5I**). CREBBP is therefore dynamically-loaded at CD40-responsive genes during the CB to CC transition in non-malignant germinal centers, and *CREBBP* KAT-PM lymphoma cells have reduced CREBBP loading at these same dynamically-loaded regions.

**Figure 5:**
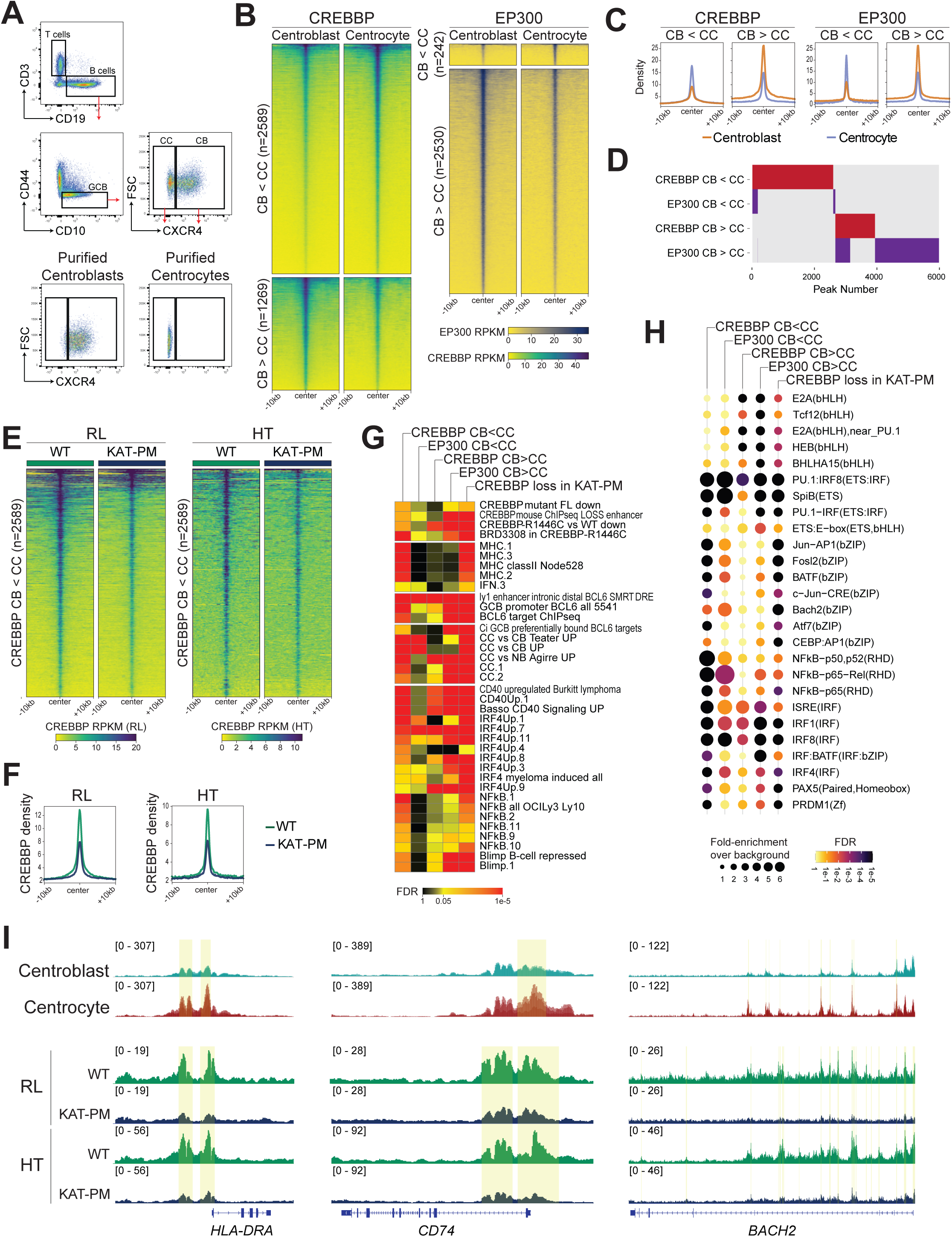
Dynamic loading of CREBBP and EP300 during centroblast to centrocyte transition. **A)** A representative example of sorting schema and purity of centroblasts and centocytes from human tonsil samples. **B-C)** Heatmaps (B) and density plots (C) show regions with significantly different loading of CREBBP (left) and EP300 (right) between centroblasts and centrocytes. **D)** Overlap of regions with differential loading of CREBBP or EP300 in centroblasts and centrocytes. **E-F)** Heatmap (E) and density plots (F) show the loading of CREBBP in isogenic WT and *CREBBP* KAT-PM lymphoma cells over regions that are dynamically by CREBBP in the centroblast to centrocyte transition. **G)** Hypergeometric gene set enrichment analysis of genes associated with peaks that are differentially loaded by CREBBP or EP300 during centroblast to centrocyte transition, or with loss of CREBBP loading in KAT-PM cells compared to isogenic WT controls. **H)** Transcription factor motif enrichment analysis of genes associated with peaks that are differentially loaded by CREBBP or EP300 during centroblast to centrocyte transition, or with loss of CREBBP loading in KAT-PM cells compared to isogenic WT controls. I) Tracks showing representative examples of regions (highlighted in yellow) with significant gain in CREBBP loading in the centroblast to centrocyte transition, and loss of CREBBP loading in KAT-PM cells compared to isogenic WT controls. Tracks represent overlays of biological replicates.

### CREBBP KAT-PM acts as a transcription factor sink

We hypothesized that the reduced loading of CREBBP KAT-PM at CD40-responsive regions and the concomitant failure of EP300 to act redundantly at these loci may be the result of CREBBP acting as a zombie enzyme (31,32). One of the mechanisms of action for inactive zombie enzymes is to inhibit the activity of its active paralog by binding shared substrates while failing to catalyze the reaction. Specifically, we predicted that CREBBP KAT-PM may inhibit EP300 by limiting quantities of transcription factor and prevent it from associating with EP300 to drive compensation. We therefore reasoned that mimicking immune synapse signaling in these cells through CD40L and IL4 treatment, which drives nuclear translocation of NFκB transcription factors and in turn activates transcription and increases the abundance of IRF4 (Figure S6A), may drive greater nuclear abundance of transcription factor and saturate CREBBP binding to allow greater association of transcription factors with EP300. We focused these experiments on the Y1503C mutant due to the slightly higher impact of catalytically-dead mutants on NFκB signatures. Differential analysis of treated and untreated WT cells identified a predominant increase in H3K27Ac (**Figure 6A-B**; 1655/2076 significant regions; FDR<0.05, fold-change>1.5; Table S11) at genes significantly for (i) those with CREBBP unloading in KAT-PM cells, (ii) those loaded by CREBBP in the CB to CC transition; (iii) signatures previously shown to be CD40-responsive and BCL6-repressed; and (iv) those regulated by IRF4 and/or NFκB (**Figure 6C**). Similar patterns were also observed for CD40/IL4-induced CREBBP-loading, supporting the notion that dynamic chromatin loading of CREBBP in centrocytes is in part responsive to immune synapse signaling.

**Figure 6:**
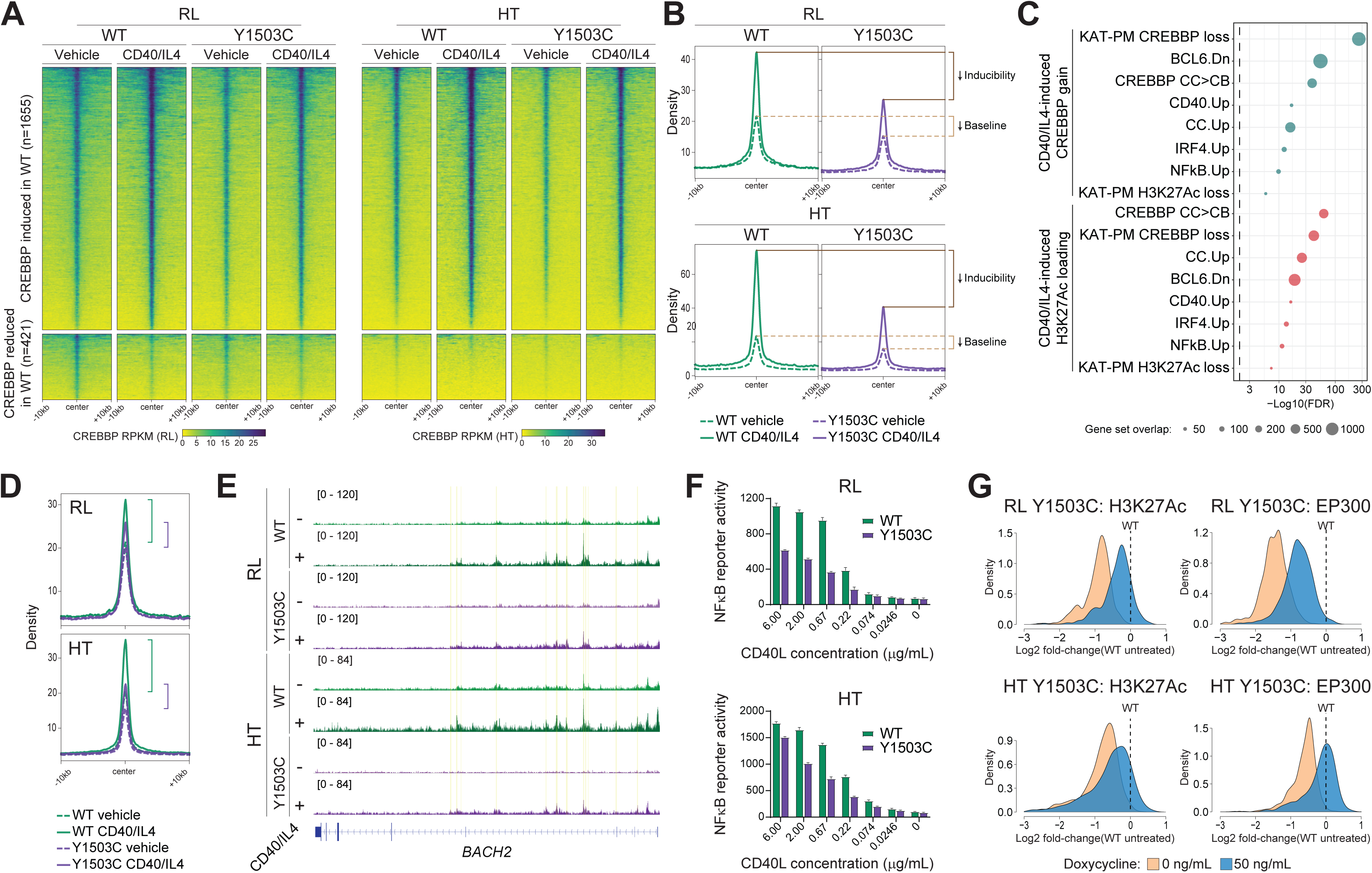
Perturbation of CD40/IL4 response in CREBBP lysine acetyltransferase domain point mutants. **A-B)** Heatmaps (A) and density plots (B) show significant (FDR≤0.05, fold-change≥1.5) changes in CREBBP loading following CD40L/IL4 treatment in isogenic WT and Y1503C mutant lymphoma cells. **C)** A bubble plot shows the size and significance of the overlap between regions with increased H3K27Ac (above) or CREBBP loading (below) following CD40L/IL4 stimulation with other gene sets defined within this study (marked by *) and prior studies. **D)** Density plots show the loading of EP300 in isogenic WT or Y1503C mutant RL (above) or HT (below) cells following CD40L/IL4 treatment. **E)** Tracks of CREBBP loading over the *BACH2* gene in isogenic WT or Y1503C mutant cells for RL (above) or HT (below) with or without CD40L/IL4 treatment. **F)** NFκB luciferase reporter activity in isogenic WT or Y1503C mutant cells treated with increasing concentrations of CD40L. **G)** Density plots, normalized to isogenic WT cells, showing the H3K27Ac (left) or EP300 loading (right) following doxycycline-inducible over-expression of the IRF4 transcription factor.

Regions with increased H3K27Ac in WT cells following CD40/IL4 treatment had markedly lower baseline levels of H3K27Ac in *CREBBP* Y1503C cells and upon failed to increase H3K27Ac levels equivalent to that observed in WT cells following treatment (**Figure 6B**). However, there was nonetheless a measurable increase in H3K27Ac that led us to evaluate EP300 loading, anchored to the regions of H3K27Ac gain in WT CD40/IL4-treated cells. We thereby observed a proportionate gain in EP300 loading at these loci in *CREBBP* Y1503C cells, showing that stimulation can at least partially overcome the deficit in EP300 loading associated with *CREBBP* KAT-PM (**Figure 6D**). These regions included genes that are important for GC exit and terminal differentiation of GC B-cells, such as multiple intronic enhancers for *BACH2* (**Figure 6E**). In order to specifically probe NFκB transcription factor activity we used a previously described NFκB luciferase reporter construct, transfected into WT and Y1403C mutant RL and HT cells, and measured response to increasing concentrations of CD40L. This showed a significant reduction in NFκB activity in Y1503C mutant cells compared to WT cells, thereby validating a blunted CD40 response in *CREBBP* mutant cells (**Figure 6F**). However, despite this reduction there remained to be a dose-dependent increase in NFκB activity with increasing concentrations of CD40L within Y1503C mutant cells (**Figure 6F**). We predict that this effect is via increased nuclear abundance of transcription factors, but CD40/IL4 signaling may have effects that extend beyond this, including potentially driving post-translational modification of CREBBP and altering transcription factor selectivity (33). We therefore engineered *CREBBP* Y1503C RL cells with transposon vector for tetracycline-inducible over-expression of IRF4 (Figure S6B) to allow for specific increase in a single transcription factor that is activated by CD40 signaling and known to interact with CREBBP (34). To achieve this, we focused specifically on peaks that satisfied all 3 of the following criteria: (i) showed increased CREBBP loading in the CB to CC transition, (ii) showed reduced CREBBP loading in KAT-PM cells, and (iii) contained one or more IRF4 motifs. Tetracycline-inducible IRF4 expression was able to increase H3K27Ac and loading of EP300 in Y1503C RL and HT cells compared to untreated controls (**Figure 6F**). Although the levels in most cases did not reach those observed at baseline in WT cells, this shows that increasing the abundance of transcription factor may be able to saturate the zombie enzyme effect of catalytically inactive CREBBP and partially restore enhancer activation via EP300 compensation.

## DISCUSSION

Catalytically inactive or lowly-active pseudoenzymes, popularly referred to as zombie enzymes, have emerged as playing an important functional role in biology by modifying the activity of their paralogs through scaffolding roles, by binding the paralog and causing allosteric changes, or by competing with the paralog for substrate without catalyzing the enzymatic reaction (31). Here we describe for the first time the role of KAT domain mutations in *CREBBP* as creating a zombie enzyme that inhibits the function of its paralog, EP300. We show by inducing increased nuclear abundance of transcription factor through CD40/IL4 stimulation or tetracycline-induced over-expression that the zombie enzyme role of CREBBP KAT-PM is likely due binding of limiting quantities of transcription factor substrate, which prevents EP300 compensation. This is further supported by the fact that knock-out of CREBBP, and a resulting absence of protein, allows for compensation by EP300 as marked by increased loading onto chromatin and maintenance of enhancer activity. The generation of a CREBBP zombie enzyme thereby constrains centrocyte enhancers that are dynamically loaded by CREBBP in the GC, restricting programs that normally regulate cell fate decisions in response to CD40 signaling. Importantly, a large number of enhancers are bound reciprocally by CREBBP and EP300 in the centroblast to centrocyte transition, with predominant chromatin-unloading of EP300 and chromatin-loading of CREBBP. Although previously considered to be largely interchangeable and redundant in many contexts, this provides important insight into the specificity of CREBBP and EP300 for different enhancer programs. These independent functions are important for normal GC B-cell development, as has been shown by compound knock-out studies in mice (8), but we expect this to extend across a range of cell types and disease states.

*CREBBP* and, to a lesser degree, *EP300* are highly recurrently mutated across cancer, including colon, breast and lung cancers (11). We suggest that KAT domain missense mutations in either of these genes are likely to confer a similarly-functioning zombie enzymes that compete with their active paralog for transcription factor binding. However, we also noted biochemical difference between classes of KAT domain mutations. While GC-derived lymphomas have a visible cluster of mutations occurring at amino acids that normally interact with the proximal portion of acetyl-CoA (acetyl group and β-mercapto-ethylamine moiety) and shape the catalytic core of the enzyme, a similar cluster of mutations is not prominently observed in either *CREBBP* or *EP300* within solid tumors. In contrast, solid tumors display a clear hostpot of mutation at the R1446 residue that we have shown normally interact with the distal portion (pyrophosphate and adenosine phosphate moieties) of acetyl-CoA. EP300 mutational hotspots in solid tumors similarly are located at amino acids that interact with the distal portion of acetyl-CoA. We have shown in our data that R1446 mutations retain some catalytic activity and are by-in-large sufficient to maintain acetylation at super-enhancers and weak enhancers. We speculate that preferential acquisition of R1446 mutations in follicular lymphoma and solid tumors may therefore be due to the ability of these residually-active mutants to maintain super-enhancer activity that might be beneficial for cell survival or oncogenesis.

Based upon our observed dynamics of CREBBP and EP300 loading and histone acetylation, underlying patterns transcription factor motif enrichment, and functional experiments modulating signaling inputs and transcription factor abundance, we propose the following overall model for signal-responsive CREBBP/EP300-mediated transcription: (i) signaling induces increased nuclear abundance of active transcription factor that is bound by CREBBP or EP300, (ii) CREBBP or EP300 associate with transcription factors in the nucleus and the complex probes accessible motifs with a low baseline residence time; (iii) local acetylation mediated by CREBBP or EP300, and a confluence of other local factors including the recruitment of bromodomain-containing proteins with intrinsically disordered domains (IDRs) may drive condensate formation that stabilize the complex on chromatin leading to increased residence time and transcriptional activation. In support of this model, the role of histone acetylation and CREBBP/EP300 function in driving transcriptional condensate formation (35), and co-condensation between CREBBP/EP300 and TFs (36), has been clearly defined. Using our CUT&RUN assays that provide a snapshot of binding across many cells at a single point in time, this appears as increased CREBBP/EP300 loading in response to stimulus. In contrast, in the setting of inactivating CREBBP mutations, local acetylation does not occur and the residence time of TF/CREBBP complexes may therefore remain low, appearing as low overall CREBBP loading when viewed as a snapshot across cells at a single time point.

In conclusion, recurrent *CREBBP* mutations differentially alter cofactor interactions leading to variable loss of activity that results in a zombie enzyme which inhibits the function of it paralog, EP300, by binding limiting amounts of TF and preventing EP300 compensation. This results in a loss developmentally-regulated dynamic loading of CREBBP that retards the activation of enhancers that are normally active within centrocytes in response to CD40 signaling. The pattern of mutations that we characterized in *CREBBP* within GC-derived lymphomas are also observed within both *CREBBP* and *EP300* in other hematological and solid cancers, suggesting that the mutational-acquisition of zombie acetyltransferase enzymes may be a novel mechanism of oncogenesis that is conserved across cancer. In other disease contexts we predict that CREBBP or EP300 may be dynamically loaded onto chromatin in response to alternative signals that regulate cell state transitions that are relevant to the respective cancer’s cell of origin, and that inhibition of this process by mutationally-encoded zombie enzymes epigenetically constrain the developmental fates of cancer cells or cancer cell precursors.

## METHODS

### Enzyme activity assay

The CREBBP acetyltransferase activity was assayed as previously described (23). Human CREBBP catalytic domain (CREBBP-KAT) (residues 1096-1751 consisting of Bromo domain, Ring domain, PHD domain and KAT domain) or mutated forms (R1446C, Y1482N, Y1503C and Y1503F) were expressed in *E. coli* and purified as described previously (37). CREBBP catalytic domain (500 nM) was incubated with 100 µM AcCoA, 30 µM H3K18 (5-FAM-RAPRK18QLAT-NH2) or H3K27 (5-TKAARKSAPAT-NH2) or p53 (5-TSRHKKLMF-NH2) peptides, and 1 mM DTT in a total volume of 100 µl of assay buffer (25 mM Tris-HCl pH 8.0) for 20 min at room temperature. The reaction was stopped using 100 µl of HClO4 (15% v/v in water) and 20 μl aliquots of the mixture were injected in a reverse phase ultra-fast liquid chromatography system (RP-UFLC, Shimadzu, France). Peptides were separated under isocratic conditions (80% acetonitrile, 20% water, 0.12% TFA, flow rate of 0.6 ml/min) on a Shim-pack XR-ODS column at 40 °C. H3K18, H3K27 and p53 substrate peptides and their acetylated product were detected by fluorescence (excitation: 485 nm, emission: 530 nm) and quantified by integration of the peak fluorescence areas

Steady state kinetics with peptide substrates (H3K18, H3K27 and p53 peptides) and Ac-CoA were carried out and catalytic efficiency determined (V_m_/K_m_) as described previously (23). Briefly, steady states assays contained CREBBP catalytic core (100 nM) with varying peptide concentrations (0-100 μM) and constant Ac-CoA (100 μM) in 50 μl activity buffer or CREBBP catalytic core (100 nM) with varying Ac-CoA concentrations (0-200 μM) and constant peptide (30 μM) in 50 μl activity buffer. CREBBP acetyltransferase activity was assayed for 20 minutes at room temperature and quantified by RP-UFLC as described above. Measurement of the acetylation of recombinant H3 and p53 proteins by CREBBP-KAT and the mutant forms were carried out as described previously using western blots with anti-H3K18ac, H3K27As, and p53ac antibodies (Zhang et al., 2021)

### Protein purification, crystallization and data collection

The pHis parallel 2 vector encoding mouse CREBBP_ΔAL (corresponding to residues 1079-1751 of mouse CREBBP with replacement of 1557-1618 by a 5-aa SGGSG linker) was used as previously reported (28). Relevant mutations were introduced with the QuikChange II XL Site-directed Mutagenesis Kit (Agilent Technologies) to generate vectors encoding R1446C/Y1503F, Y1482N, Y1503C and Y1503F CREBBP_ΔAL proteins. The different vectors were transformed into competent *E.coli* BL21(DE3) which were grown in 2×YT medium containing 30 μg/ml kanamycin and 20 μg/ml chloramphenicol. Production of the recombinant proteins was induced by 1 mM IPTG at 15 °C overnight. Bacteria pellets were resuspended in buffer A (20 mM Tris-HCl, 300 mM NaCl, 25 μM ZnSO4, 1% Triton X-100, 2 mM DTT, 1 mg/ml lysozyme, EDTA-free protease inhibitor mixture (Roche), 250 units of benzonase (Novagen), pH 8.0) and lysed for 30 min at 4°C. After sonification, the soluble fraction (obtained by centrifugation with 13000g at 4°C for 30 min) was applied to His-Select Nickel Affinity Gel (Sigma-Aldrich) in presence of 5 mM imidazole for 2h on ice. After extensive washes with buffer A, proteins were eluted with buffer B (20 mM Tris-HCl, 300 mM NaCl, 25 μM ZnSO4, pH 8.0) containing 300 mM imidazole and buffer exchanged to buffer C (Tris-HCl 25 mM, NaCl 150 mM, 25 μM ZnSO4, pH 8.0). 6xHis tag was removed by TEV with 1 mM DTT at 8 °C for overnight and the different CREBBP_ΔAL proteins were further purified by gel filtration in buffer D (Tris-HCl 25 mM, NaCl 150 mM, 25μM ZnSO4, 2 mM DTT, pH 8.0). The purified proteins 10 mg/ml were mixed with CoA at a ratio of 1:10 in buffer D and used for crystallization screening using the sitting drop vapour diffusion method with the Mosquito (TTP Labtech) automated crystallization system. 672 different crystallization solutions were mixed with prepared proteins. Crystals were reproduced manually according to hit conditions using hanging drop vapour-diffusion technique (at 18°C) and grown by mixing protein and reservoir solution containing 0,2 M Sodium citrate tribasic dihydrate, 20 % w/v Polyethylene glycol 3,350 with pH 8.2 at 18 °C in different volume ratio, including 1:1, 1:2, 2:1. Crystals were then flash-cooled in liquid nitrogen using reservoir solution supplemented with 25% (vol/vol) ethylene glycol as cryoprotectant. X-ray diffraction data were collected on beamline PROXIMA-1 at the SOLEIL synchrotron (St. Aubin, France). All CREBBP_ΔAL protein structures were determined by molecular replacement method with CCP4/PHASER using CREBBP WT structure (PDBID 5U7G) as searching model. The CoA conformation was determined by PHENIX/LigandFit. Numbers of iterations of structure building and refinement were carried out by COOT, using local NCS restraints on atom position.

### Transfection of HEK 293T cells

HEK 293T cells were cultured in an incubator at 37 °C, 5% CO_2_ in DMEM medium containing 10% fetal calf serum (FCS) and 1 mM L-glutamine. Cells (3×10^6^ cells) were seeded in a 10 cm dish and transfected with 6 μg of FLAG-tagged human CREBBP plasmids (Origene,USA) using Metafectene (Biontex, Germany). After 48 h transfection, cells were lysed and supernatant was used for immunoprecipitation and western blot. Cell pellet was used for histones extraction and western blot.

### Cell extracts and histone extraction

Cells were lysed for 30 minutes at 4 °C with a lysis buffer (PBS pH 7.5, 0.5% Triton X-100, 3.3 µM TSA, cocktail protease inhibitor) and centrifuged for 10 minutes at 15000 g. The supernatant (cytoplasmic fraction) was harvested and the pellet (nucleus enriched fraction) was incubated overnight at 4 °C with 400 μl of 0.2N HCl. The next day the acid extract was centrifuged (10 minutes at 6500 g) and the supernatant (fraction enriched in histones) saved. Histones (2 μg) were separated by 18% SDS-PAGE and analyzed by western blot using H3K18Ac, H3K27Ac or H3 antibodies (Cell signaling).

### Immunoprecipitation

The cells were lysed with lysis buffer (PBS pH 7.5, 1% Triton X-100, 3.3 µM TSA, 100 μM DTT, protease inhibitor cocktail) for 30 min on ice and centrifuged at 15000 g for 15 min at 4 °C. The protein concentration of cell extracts was determined by the Bradford method.

Immunoprecipitation was carried out overnight at 4 °C with 1 μg of anti-CREBBP antibody (A22, Santa Cruz) or 1 μg of anti-flag (α-DDK, TA50011, Origene) in 1 ml of lysis buffer containing 1 mg of total protein extract. The following day, 25 μl of Protein A or G agarose beads (H0116, Santa cruz) were added and the mixture incubated for 2 hours at 4 °C. The beads were washed 3 times with assay buffer and then incubated at room temperature with 60 μl of assay buffer (25 mM Tris-HCl pH 8.0). After 3 hours of incubation, the beads were sedimented by centrifugation and an aliquot of 50 µl of supernatant was taken and mixed with 50 µl of HClO4 was added. The activity immunoprecipitated CREBBP proteins was determined by UFLC as previously above. The beads were washed with PBS and then incubated with Laemmli buffer containing 200 mM β-ME for 5 min at 100 °C. Then 20 µl of supernatant was used for SDS-PAGE and western blot.

### Western blot

Samples were separated by SDS-PAGE prior to being transferred onto a nitrocellulose membrane (0.45µ, GE Healthcare) for 2 h at 4°C. Membranes were blocked in 5% non-fat milk PBST for 2h and incubated with primary antibody in 1% non-fat milk PBST at 4 °C for overnight under agitation. The next day, membranes were washed 5 times with PBST and incubated with secondary antibody for 2 h at room temperature under agitation. Membranes were then washed 5 more times and the signal was detected by chemiluminescence using ECL Prime (GE Healthcare) on an Amersham Imager 600 detection system (GE Healthcare).

### Thermal shift assay (TSA)

Thermal shift assays were performed to measure the thermal stability of CREBBP ΔAL wild-type or mutant proteins alone or in the presence of ligands. CREBBP ΔAL (1.5 μM) was incubated in the presence of 10X SYPRO orange dye (S5692, Sigma), 400 μM CoA and/or 1 mM H3K18 peptide (GGKAPRKQLATKAA) in a total volume of 10 μl buffer E (25mM Tris, pH 8.0, 300mM NaCl) for 10 min at room temperature in 384-well opaque (Roche) PCR plates. Plates were then centrifuged (1000 g, 4°C) and real-time analysis of the thermal stability of the protein was performed by fluorescence reading (excitation: 465 nm, emission: 580 nm) on a Light Cycler 480 (Roche).

### Statistics, protein structure and image analysis

Values are presented as mean ± standard deviation (SD) and analyzed by oneway analysis of variance (ANOVA) using GraphPad Prism 5. Grayscale quantitative analysis was performed using Image Pro-Plus 6.0. Crystal structures analyzed and visualized with Pymol 2.5 or UCSF Chimera x.

### Cell culture, CRISPR/Cas9 editing and validation

RL (ATCC CRL-2261), HT (ATCC CRL-2260) and HEK-293T (ATCC CRL-3216) were purchased from American Type Culture Collection. Both parental cell lines and CRISPR-modified cell lines were cultured in RPMI medium (for RL and HT) or DMEM medium (for HEK-293T) with 10% FBS and 1X penicillin/streptomycin (Corning 30-002-CI). IRF4 tetracycline-inducible cell lines were cultured in RPMI medium with 10% tetracycline negative FBS (Corning 35-075-CV). The CREBBP KAT domain mutations were introduced by Alt-R CRISPR-Cas9 (Integrated DNA Technologies) electroporation with SpCas9 HiFi nuclease (Integrated DNA Technologies 1081061), ATTO550 (Integrated DNA Technologies 1075928) labelled guide RNA (Table S12) and a single-stranded oligonucleotide donor template (Table S12). Single ATTO550-positive cells were sorted into 96-well plate after 48 hours of electroporation and expanded followed by validation using Sanger sequencing of the amplicon from the targeted regions (Table S12). All cells were maintained as sub-confluent culture and mycoplasma tested prior to each set of assays.

### CUT&RUN

CUT&RUN was performed as previously described (29,38,39), with some modifications. Cells were collected and washed by PBS. After final wash, nuclei were isolated using nuclei extraction buffer (Table S13) and then washed twice with wash buffer (Table S13). Five-hundred thousand nuclei were counted and incubated with activated Concanavalin A coated beads (EpiCypher 21-1411) and subsequently mixed with antibody (Table S14) in antibody buffer (Table S13) for overnight incubation on nutator in a cold room. IgG controls were set in parrel for antibody enrichment and specificity validation. Nuclei were then washed three times with digitonin buffer (Table S13) and incubated with PAG-MNase (EpiCypher 15-1116) for 10 minutes at room temperature. After 10 minutes of incubation, nuclei were washed three times again with digitonin buffer and resuspended with prechilled low salt high Ca2+ buffer (Table S13) to initiate 1-hour digestion at 0 °C (ice block). The reaction was terminated by switching into stop buffer (Table S13) and incubating at 37 °C for 10 minutes to release digested DNA from the nuclei. DNA was then purified with Ampure beads (Beckman A63882) size selection (0.5X to remove large fragment and 2X to recover desired nucleosomal DNA fragments). NGS libraries were generated with KAPA Hyper Prep Kits (Roche KK8502) using 14 cycles of PCR amplification. Libraires were validated on a Tapestation 4200 (Agilent G2991BA), quantified by Qubit High Sensitivity dsDNA Kit (Life Technologies Q32854), multiplexed and sequenced on a NovaSeq6000 using 100-10-10-100bp reads format.

### ATAC-sequencing

ATAC-seq was performed as previously described (40). Fifty thousand cells were washed in ATAC-seq resuspension buffer (Table S13) were resuspended in 50 μL of in ATAC-seq resuspension buffer containing 0.1% NP-40, 0.1% Tween-20 and 0.01% digitonin, and incubated on ice for 3 min. After lysis, nuclei were wash with cold ATAC-seq resuspension buffer containing 0.1% Tween-20 but no NP40 or digitonin, subsequently resuspended in 50 μL of transposition mixture (Table S13) and incubated at 37 °C for 30 minutes in a thermomixer with 1000 RPM shaking. Fragmented DNA were then extracted with Zymo DNA Clean and Concentrator-5 kit (Zymo D4024). NGS libraries were prepared with KAPA Hyper Prep Kits (Roche KK8502) using 10 cycles of PCR amplification. Libraires were validated on a Tapestation 4200 (Agilent G2991BA), quantified by Qubit High Sensitivity dsDNA Kit (Life Technologies Q32854), multiplexed and sequenced on a NovaSeq6000 using 100-10-10-100bp reads format.

### RNA-sequencing

RNA-seq was performed as previously described (9). Total RNA was extracted using AllPrep DNA/RNA Kits (Qiagen 80204) and quantified with Qubit High Sensitivity RNA Kit (Life Technologies Q32852). Total RNA (1 μg) was used to generate RNA-seq library with KAPA HyperPrep RNA Kits with RiboErase (Roche KK8560) using 8 cycles of PCR amplification. Libraires were validated on a Tapestation 4200 (Agilent G2991BA), quantified by Qubit High Sensitivity dsDNA Kit (Life Technologies Q32854), multiplexed and sequenced on a NovaSeq6000 using 100-10-10-100bp reads format.

### Isolation of B cell populations from human tonsil by FACS

Primary cells were isolated from fresh deidentified human tonsillar lymph nodes by mechanical tissue dissociation and cryopreserved in 90% FBS with 10% DMSO. Prior to each profiling, cells were revived and resuspended in RPMI medium with 10% FBS at 37 °C for 1 hour. After washing with PBS, cells were labelled with anti-CD3 (1:20 dilution), anti-CD19 (1:20 dilution), anti-CD10 (1:20 dilution), anti-CD44 (1:20 dilution) and anti-CXCR4 (1:20 dilution) conjugated fluorochromes for 30 minutes, followed by 3 times of washing in PBS with 2% BSA. Cells were sorted on a BD FACSAria™ Fusion cytometer from the Advanced Cytometry & Sorting Facility at MD Anderson Cancer Center. Post-sorting analysis were performed immediately after each sorting to ensure the purity of each population. Antibody details can be found in Table S13.

### CD40 stimulation

CD40 stimulation medium were prepared by incubation of 40 uL of CD40 ligand and 40 uL of cross-linking antibody (Miltenyi Biotec 130-106-196) mixture at room temperature for 30 minutes and subsequently adding into 10 mL of RPMI+10%FBS and 2 uL of IL4 (Miltenyi Biotec 130-106-196). Cells were seeded at 0.5 × 10^6^ per mL of CD40 stimulation medium at 37 °C. Negative control group was set in the absence of CD40 ligand multimer and IL4. For CUT&RUN, cells were collected after 24 hours of stimulation. For immunoblotting, cells were collected after 24 hours of stimulation and performed with cell fractionation using Nuclear Extract Kit (Active Motif 40410). For flow cytometry analysis, cells were collected after 1, 3 and 7 days, followed by antibody staining and flow cytometry testing.

### Viral transduction and NF-κB reporter assay

Lentivirus was produced in 293T cells by co-transfecting NF-κB reporter (41) with packaging vectors psPAX2 (Addgene #12260, a gift from Didier Trono) and pMD2.G (Addgene #12259, a gift from Didier Trono) at a 12:10:5 ratio in OptiMEM (Thermo Fisher 31985070) with Lipofectamine 2000 transfection reagent (Thermo Fisher 11668019). Virus particles were collected from the supernatant 48 hours after transfection, filtered through a 0.45-μm filter, and mixed with polybrene infection reagent (Sigma-Aldrich TR-1003-G) prior to the 2-hour spinfection with lymphoma cell lines. Cells were selected 24 hours after infection by adding puromycin (Invivogen ant-pr-1) at 2ug/mL concentration for a week. Stable cell lines with NF-κB reporter were treated with difference concentration of CD40L and IL2 for 24 hours, followed by measuring luciferase activities using ONE-Glo™ EX Luciferase Assay System (Promega E8110) according to the manufacturer’s instructions. All assays were repeated with three independent experiments.

### Cloning and tetracycline inducible expression of IRF4

The IRF4 gene was cloned into pSBtet-GP backbone (Addgene #60495, a gift from Eric. Kowarz) by replacing luciferase gene with IRF4 Human Tagged ORF (Origene #RC204876). IRF4 tetracycline-inducible cell lines (RL CREBBP^WT^, RL CREBBP^Y1503C^, HT CREBBP^WT^ and HT CREBBP^Y1503C^) were generated by co-electroporation of transposase-expressing vector pCMV(CAT)T7-SB100 (Addgene #34879, a gift from Zsuzsanna. Izsvak) and pSBtet-GP vector that expresses IRF4 using neon transfection system (Invitrogen MPK10096). Transfected cells were further purified by puromycin (1 μg/mL) selection for 10 days and maintained in RPMI medium with 10% tetracycline negative FBS (Corning 35-075-CV). Stable cell lines were validated by immunoblotting after a dose titration of doxycycline (Thermo Fisher J60579.22) induction for 72 hours (Figure S6B). Cell lines were seeded at 0.25 × 10^6^ per mL and induced with 50 ng doxycycline per mL of medium for 72 hours prior to CUT&RUN assay.

### Immunoblotting

Whole-cell lysates were extracted with RIPA buffer and quantified by DC Protein Assay (Biorad 5000111). Protein samples were separated by SDS-PAGE gels (Biorad 4561086) and transferred to PVDF membranes. After blocking with 5% non-fat dry milk for 1 hour, membranes were incubated with the indicated primary antibodies: anti-CREBBP (dilution 1:1000), anti-EP300 (dilution 1:1000), anti-p50/p105 (dilution 1:1000), anti-p65 (dilution 1:2000), anti-IRF4 (dilution 1:1000), anti-CIITA (dilution 1:1000), anti-GAPDH (dilution 1:2000), anti-β-Actin (dilution 1:2000), and then anti-mouse/rabbit peroxidase-conjugated secondary antibodies (dilution 1:2000). Protein signal was detected with enhanced chemiluminescence using the Bio Rad ChemiDoc imaging system. Antibody details can be found in Table S13.

### SNAP-ChIP qPCR

All histone markers antibodies for CUT&RUN were either certified by EpiCypher (H3K4me1, H3K4me3, H3K27me3) or validated by SNAP-ChIP qPCR (H3K27Ac). To validate the specificity and target enrichment of H3K27Ac antibody for CUT&RUN, H3K27Ac antibody was incubated with SNAP-ChIP® K-AcylStat™ Panel (EpiCypher 19-3100) according to the manufacturer’s instructions, followed by immunoprecipitation with Dynabeads™ Protein G (Thermo Fisher 10004D). Input control was reserved without adding antibody. Immunoprecipitated DNA was purified by 2X Ampure beads (Beckman A63882) and utilized for qPCR validation with SNAP-ChIP K-AcylStat qPCR primers (Table S12).

### Data processing and statistical analysis

#### CUT&RUN processing

CUT&RUN processing was performed as previously described (42), with some modifications. Sequencing quality control was evaluated by FastQC. Paired-end CUT&RUN raw reads were trimmed to remove adapter sequences using Trimmomatic and then aligned to version 19 of the human reference genome (hg19) using Bowtie2 with options: --local --very-sensitive --no-mixed --no-discordant --phred33 -I 10 -X 700. Aligned reads were sorted by coordinates and tested for mapped fragment size distribution by Samtools. The final aligned and sorted Bam files were used for further processing. Visualization was assessed by converting the final Bam files to Bigwig files using the ‘bamCoverage’ command in deepTools with option: --normalizeUsing RPKM -e -bs 50 and evaluated in IGV browser (Broad Institute). CUT&RUN peaks were called by MACS2 callpeak with option: -g hs -B -f BAM -q 0.01. IgG controls were used for comparison. Each technical repeat and biological repeat were separately processed.

#### ATAC-seq processing

Technical duplicates of each ATAC-seq samples were combined and sequencing quality control was evaluated by FastQC. Fastq files of paired-end ATAC-seq data were trimmed to remove adapter sequences using Trimmomatic and then aligned to version 19 of the human reference genome (hg19) using BWA (Burrows-Wheeler Aligner). Aligned reads were sorted by coordinates and filtered out mitochondrial DNA and black-listed regions by Samtools. Duplicate read pairs were marked and removed using the ‘MarkDuplicates’ command in picard tools. The final aligned, sorted and filtered Bam files were used for further analysis. ATAC-seq visualization was assessed by converting the final Bam files to Bigwig files using the ‘bamCoverage’ command in deepTools and corrected for Tn5 insertion. ATAC-seq peaks were called from Tn-5 corrected insertions using MACS2 callpeak with option ‘-g hs -q 0.05 -f BAMPE and -B’.

#### RNA-seq analysis

RNA-seq data was trimmed with Trimmomatic, and both pre-and post-trimming quality controls were evaluated by FastQC. The trimmed data were aligned to version 19 of the human reference genome (hg19) using STAR (2-pass). Samples with at least 20 million uniquely mapping reads were retained for further analysis. Reads count was performed with the ‘Htseq-count’ command in HTSeq with option: -m union -f bam -i gene_id -s no -t exon --additional-attr=gene_name. Reads count filtering was applied to the DESeqDataSet matrix to keep only rows that have at least 10 reads total. Differential gene expression analysis was performed using DESeq2 package in R. Principal components were calculated by DESeq2 and plotted by ggplot2.

### Identification of super enhancers

H3K27Ac peaks called from CUT&RUN were used to define the enhancer boundary, followed by further filtering based on the criteria: (1) excluding H3K27Ac peaks that overlapped with H3K4me3 peaks; and (2) excluding H3K27Ac peaks that overlap with known annotated gene promoters (±1.5 kb from the TSS). The super enhancers were identified by using the ROSE (Rank Ordering of Super-Enhancers) algorithm (30) based on the enrichment profile as the inflection point of the difference between the H3K27Ac CUT&RUN signal and IgG CUT&RUN signal versus the rank of stitched enhancers with the following parameters: -s 13500 -t 1500.

### Consensus peak and differential peak calling

To ensure the reproducibility of sample replicates, peaks called from each biological replicates and/or technical replicates were merged in Diffbind package in R and consensus peaks were determined by calculating their RPKM scores from final aligned and sorted Bam files and filtering out intervals with RPKM less than 1 across all replicates. Differential analyses were further performed using command dba.count(dbObj, bUseSummarizeOverlaps=TRUE, score=DBA_SCORE_RPKM, summits=FALSE) from DiffBind with consensus peaks or defined peak sets as input. RPKM values were calculated by dba.peakset(dbObj, bRetrieve=TRUE) for plotting log2fold change histogram and identifying principal components and their coordinates. Differential regions were determined by command dba.report(dbObj, method=DBA_DESEQ2, contrast = 1, th = 1). The statistical thresholds for significance were FDR < 0.05 and fold change > 1.5.

### Definition and visualization of peak sets

H3K27Ac consensus peaks defined in wild-type cell lines were applied as input peak set for DiffBind to determine the differential peak sets of H3K27Ac loss in R1446C and Y1482N/Y1503C mutants. The intersection of two differential peak sets were defined as the shared loss regions using Bedtools intersect. The complement of two peak sets were defined as the unique loss regions using Bedtools subtract. To determine the peak sets of super enhancers, typical enhancers, weak enhancers, and active promoters, H3K27Ac consensus peaks were dissected on the basis of the ranking by ROSE algorithm (super enhancer), regions in the presence of H3K4me1 only but not in super enhancers (typical enhancers), regions in the presence of H3K4me3 only (active promoters), and regions in the absence of both H3K4me1 and H3K4me3 (weak enhancers). CREBBP/EP300 co-bound H3K27Ac loss regions in KAT domain point mutants were defined by the consensus regions of H3K27Ac loss between wide-type and all three mutants (R1446C, Y1482N and Y1503C), using the overlap regions of H3K27Ac, CREBBP and EP300 as input peak set for DiffBind. Differential peak sets of CREBBP/EP300 between centroblasts and centrocytes were computed using DiffBind. The overlaps and differences of dynamically loaded regions were determined by Bedtools intersect and subtract respectively, and visualized by summarizing the counts of their intersection and complement in ggplot2 package in R. All heatmap and profile plots of CUT&RUN were plotted by deepTools with command ‘computeMatrix’, ‘plotHeatmap’ and ‘plotProfile’.

### Hypergeometric gene set enrichment analysis (hGSEA)

Differential peaks were first annotated to the genes with the closest transcription start site to each peak by HOMER (UCSD) annotatePeaks.pl. Hypergeometric gene set enrichment analysis (hGSEA) was performed using hypeR package in R for differential genes, with our curated normal B cell and B cell lymphoma gene sets and the SignatureDB gene set database (https://lymphochip.nih.gov/signaturedb/). The statistical threshold for significance of hGSEA was FDR < 0.05.

### Motif enrichment

Motif enrichment analysis was performed using HOMER (UCSD) findMotifsGenome.pl, with previously identified differential peaks as input, to check the enrichment of known motifs in HOMER Motif Databases (http://homer.ucsd.edu/homer/motif/HomerMotifDB/homerResults.html). The option “-size 200” was set to determine the size of the region for motif searching. Background regions were chosen randomly by Homer based on normalized CpG% or GC% content. The statistical threshold for significance of motif enrichment was FDR < 0.05.

## REFERENCES

1. Berdasco M, Esteller M. Aberrant epigenetic landscape in cancer: how cellular identity goes awry. Dev Cell 2010;19(5):698–711 doi 10.1016/j.devcel.2010.10.005.

2. Feinberg AP, Levchenko A. Epigenetics as a mediator of plasticity in cancer. Science 2023;379(6632):eaaw3835 doi 10.1126/science.aaw3835.

3. Lunning MA, Green MR. Mutation of chromatin modifiers; an emerging hallmark of germinal center B-cell lymphomas. Blood cancer journal 2015;5:e361 doi 10.1038/bcj.2015.89.

4. De Silva NS, Klein U. Dynamics of B cells in germinal centres. Nature reviews Immunology 2015;15(3):137–48 doi 10.1038/nri3804.

5. Green MR. Chromatin modifying gene mutations in follicular lymphoma. Blood 2018;131(6):595–604 doi 10.1182/blood-2017-08-737361.

6. Pasqualucci L, Dominguez-Sola D, Chiarenza A, Fabbri G, Grunn A, Trifonov V, et al. Inactivating mutations of acetyltransferase genes in B-cell lymphoma. Nature 2011;471(7337):189-95 doi 10.1038/nature09730.

7. Mullighan CG, Zhang J, Kasper LH, Lerach S, Payne-Turner D, Phillips LA, et al. CREBBP mutations in relapsed acute lymphoblastic leukaemia. Nature 2011;471(7337):235-9 doi 10.1038/nature09727.

8. Meyer SN, Scuoppo C, Vlasevska S, Bal E, Holmes AB, Holloman M, et al. Unique and Shared Epigenetic Programs of the CREBBP and EP300 Acetyltransferases in Germinal Center B Cells Reveal Targetable Dependencies in Lymphoma. Immunity 2019;51(3):535–47 doi 10.1016/j.immuni.2019.08.006.

9. Mondello P, Tadros S, Teater M, Fontan L, Chang AY, Jain N, et al. Selective Inhibition of HDAC3 Targets Synthetic Vulnerabilities and Activates Immune Surveillance in Lymphoma. Cancer discovery 2020;10(3):440–59 doi 10.1158/2159-8290.CD-19-0116.

10. Tate JG, Bamford S, Jubb HC, Sondka Z, Beare DM, Bindal N, et al. COSMIC: the Catalogue Of Somatic Mutations In Cancer. Nucleic acids research 2019;47(D1):D941–D7 doi 10.1093/nar/gky1015.

11. Consortium ITP-CAoWG. Pan-cancer analysis of whole genomes. Nature 2020;578(7793):82-93 doi 10.1038/s41586-020-1969-6.

12. Green MR, Kihira S, Liu CL, Nair RV, Salari R, Gentles AJ, et al. Mutations in early follicular lymphoma progenitors are associated with suppressed antigen presentation. Proc Natl Acad Sci U S A 2015;112(10):E1116–25 doi 10.1073/pnas.1501199112.

13. Ma MCJ, Tadros S, Bouska A, Heavican T, Yang H, Deng Q, et al. Subtype-specific and co-occurring genetic alterations in B-cell non-Hodgkin lymphoma. Haematologica 2021;107(3):690–701 doi 10.3324/haematol.2020.274258.

14. Kridel R, Chan FC, Mottok A, Boyle M, Farinha P, Tan K, et al. Histological Transformation and Progression in Follicular Lymphoma: A Clonal Evolution Study. PLoS Med 2016;13(12):e1002197 doi 10.1371/journal.pmed.1002197.

15. Krysiak K, Gomez F, White BS, Matlock M, Miller CA, Trani L, et al. Recurrent somatic mutations affecting B-cell receptor signaling pathway genes in follicular lymphoma. Blood 2017;129(4):473–83 doi 10.1182/blood-2016-07-729954.

16. Jurinovic V, Kridel R, Staiger AM, Szczepanowski M, Horn H, Dreyling MH, et al. Clinicogenetic risk models predict early progression of follicular lymphoma after first-line immunochemotherapy. Blood 2016;128(8):1112–20 doi 10.1182/blood-2016-05-717355.

17. Bouska A, McKeithan TW, Deffenbacher KE, Lachel C, Wright GW, Iqbal J, et al. Genome-wide copy-number analyses reveal genomic abnormalities involved in transformation of follicular lymphoma. Blood 2014;123(11):1681–90 doi 10.1182/blood-2013-05-500595.

18. Reddy A, Zhang J, Davis NS, Moffitt AB, Love CL, Waldrop A, et al. Genetic and Functional Drivers of Diffuse Large B Cell Lymphoma. Cell 2017;171(2):481–94 e15 doi 10.1016/j.cell.2017.09.027.

19. Chapuy B, Stewart C, Dunford AJ, Kim J, Kamburov A, Redd RA, et al. Molecular subtypes of diffuse large B cell lymphoma are associated with distinct pathogenic mechanisms and outcomes. Nature medicine 2018;24(5):679–90 doi 10.1038/s41591-018-0016-8.

20. Schmitz R, Wright GW, Huang DW, Johnson CA, Phelan JD, Wang JQ, et al. Genetics and Pathogenesis of Diffuse Large B-Cell Lymphoma. The New England journal of medicine 2018;378(15):1396–407 doi 10.1056/NEJMoa1801445.

21. Lacy SE, Barrans SL, Beer PA, Painter D, Smith AG, Roman E, et al. Targeted sequencing in DLBCL, molecular subtypes, and outcomes: a Haematological Malignancy Research Network report. Blood 2020;135(20):1759–71 doi 10.1182/blood.2019003535.

22. Karube K, Enjuanes A, Dlouhy I, Jares P, Martin-Garcia D, Nadeu F, et al. Integrating genomic alterations in diffuse large B-cell lymphoma identifies new relevant pathways and potential therapeutic targets. Leukemia 2018;32(3):675–84 doi 10.1038/leu.2017.251.

23. Duval R, Fritsch L, Bui LC, Berthelet J, Guidez F, Mathieu C, et al. An acetyltransferase assay for CREB-binding protein based on reverse phase-ultra-fast liquid chromatography of fluorescent histone H3 peptides. Analytical biochemistry 2015;486:35–7 doi 10.1016/j.ab.2015.06.024.

24. Henry RA, Kuo YM, Andrews AJ. Differences in specificity and selectivity between CBP and p300 acetylation of histone H3 and H3/H4. Biochemistry 2013;52(34):5746–59 doi 10.1021/bi400684q.

25. Thompson PR, Wang D, Wang L, Fulco M, Pediconi N, Zhang D, et al. Regulation of the p300 HAT domain via a novel activation loop. Nature structural & molecular biology 2004;11(4):308–15 doi 10.1038/nsmb740.

26. Liu X, Wang L, Zhao K, Thompson PR, Hwang Y, Marmorstein R, et al. The structural basis of protein acetylation by the p300/CBP transcriptional coactivator. Nature 2008;451(7180):846-50 doi 10.1038/nature06546.

27. Kaczmarska Z, Ortega E, Goudarzi A, Huang H, Kim S, Marquez JA, et al. Structure of p300 in complex with acyl-CoA variants. Nature chemical biology 2017;13(1):21–9 doi 10.1038/nchembio.2217.

28. Park S, Stanfield RL, Martinez-Yamout MA, Dyson HJ, Wilson IA, Wright PE. Role of the CBP catalytic core in intramolecular SUMOylation and control of histone H3 acetylation. Proc Natl Acad Sci U S A 2017;114(27):E5335–E42 doi 10.1073/pnas.1703105114.

29. Skene PJ, Henikoff S. An efficient targeted nuclease strategy for high-resolution mapping of DNA binding sites. Elife 2017;6 doi 10.7554/eLife.21856.

30. Whyte WA, Orlando DA, Hnisz D, Abraham BJ, Lin CY, Kagey MH, et al. Master transcription factors and mediator establish super-enhancers at key cell identity genes. Cell 2013;153(2):307–19 doi 10.1016/j.cell.2013.03.035.

31. Murphy JM, Farhan H, Eyers PA. Bio-Zombie: the rise of pseudoenzymes in biology. Biochem Soc Trans 2017;45(2):537–44 doi 10.1042/BST20160400.

32. Eyers PA, Murphy JM. The evolving world of pseudoenzymes: proteins, prejudice and zombies. BMC Biol 2016;14(1):98 doi 10.1186/s12915-016-0322-x.

33. Huang WC, Ju TK, Hung MC, Chen CC. Phosphorylation of CBP by IKKalpha promotes cell growth by switching the binding preference of CBP from p53 to NF-kappaB. Molecular cell 2007;26(1):75–87 doi 10.1016/j.molcel.2007.02.019.

34. Conery AR, Centore RC, Neiss A, Keller PJ, Joshi S, Spillane KL, et al. Bromodomain inhibition of the transcriptional coactivators CBP/EP300 as a therapeutic strategy to target the IRF4 network in multiple myeloma. Elife 2016;5 doi 10.7554/eLife.10483.

35. Gibson BA, Doolittle LK, Schneider MWG, Jensen LE, Gamarra N, Henry L, et al. Organization of Chromatin by Intrinsic and Regulated Phase Separation. Cell 2019;179(2):470–84 e21 doi 10.1016/j.cell.2019.08.037.

36. Ma L, Gao Z, Wu J, Zhong B, Xie Y, Huang W, et al. Co-condensation between transcription factor and coactivator p300 modulates transcriptional bursting kinetics. Molecular cell 2021;81(8):1682–97 e7 doi 10.1016/j.molcel.2021.01.031.

37. Zhang W, Berthelet J, Michail C, Bui LC, Gou P, Liu R, et al. Human CREBBP acetyltransferase is impaired by etoposide quinone, an oxidative and leukemogenic metabolite of the anticancer drug etoposide through modification of redox-sensitive zinc-finger cysteine residues. Free Radic Biol Med 2021;162:27–37 doi 10.1016/j.freeradbiomed.2020.11.027.

38. Skene PJ, Henikoff JG, Henikoff S. Targeted in situ genome-wide profiling with high efficiency for low cell numbers. Nature protocols 2018;13(5):1006–19 doi 10.1038/nprot.2018.015.

39. Meers MP, Bryson TD, Henikoff JG, Henikoff S. Improved CUT&RUN chromatin profiling tools. Elife 2019;8 doi 10.7554/eLife.46314.

40. Corces MR, Trevino AE, Hamilton EG, Greenside PG, Sinnott-Armstrong NA, Vesuna S, et al. An improved ATAC-seq protocol reduces background and enables interrogation of frozen tissues. Nature methods 2017;14(10):959–62 doi 10.1038/nmeth.4396.

41. Xia M, David L, Teater M, Gutierrez J, Wang X, Meydan C, et al. BCL10 Mutations Define Distinct Dependencies Guiding Precision Therapy for DLBCL. Cancer discovery 2022;12(8):1922–41 doi 10.1158/2159-8290.CD-21-1566.

42. Henikoff S, Henikoff JG, Kaya-Okur HS, Ahmad K. Efficient chromatin accessibility mapping in situ by nucleosome-tethered tagmentation. Elife 2020;9 doi 10.7554/eLife.63274.

